# Limited evolution despite years of measurable viremia in a cART-treated seronegative HIV-1 positive individual

**DOI:** 10.1101/2020.02.20.957274

**Authors:** Helen R. Fryer, Jayna Raghwani, M John Gill, Guido van Marle, Tanya Golbchik, Joe Grove, Katrina A. Lythgoe

## Abstract

Understanding the role that antibodies play in controlling HIV-1 infection and in the dynamics that underpin the formation of the HIV-1 reservoir are important steps towards combatting this global disease. To address these gaps, we performed whole-genome, deep sequence analysis of longitudinal plasma HIV-1 samples from an individual who failed to develop detectable anti-HIV-1 antibodies for 4 years post infection. These analyses reveal limited evolution despite months of measurable viremia during treatment with cART. We used a mathematical model to simultaneously analyse the viral and evolutionary dynamics of this unique individual. We propose a role for antibodies in reducing viral infectivity and demonstrate how our data are consistent with a theory of rapid activation of latently infected cells prior to effective viral suppression. Our study supports and elucidates a recent finding that although the latent reservoir persists for years once virus is effectively suppressed, prior to suppression, viral strains within the reservoir turn over rapidly. The implications for a cure are significant.

## Introduction

Following human immunodeficiency virus-1 (HIV-1) infection, anti-HIV-1 antibodies usually become detectable from approximately 2-3 weeks after virus acquisition [1]. Although some individuals are slower to seroconvert, a handful of individuals have been identified who remain persistently (i.e. greater than 4 months) seronegative. An individual who experienced the longest identified delay between acute HIV-1 infection and seroconversion – 48 months – has previously been investigated [2, 3]. Here we combine a mathematical framework with deep-sequence analysis to examine the virus and evolutionary dynamics of this unique individual. By comparing these dynamics to those observed in typical, seropositive HIV-1 individuals [4–10], this study provides a rare glimpse into the role that antibodies play in controlling HIV-1 infection. Furthermore, it allows us to explore the dynamics that underpin the formation and maintenance of the HIV-1 reservoir.

Understanding the role that antibodies play in controlling natural HIV-1 infection is vital for the design of both an HIV-1 vaccine and a cure. Although antibodies capable of neutralizing virions can typically be identified during HIV-1 infection [11], most have impaired efficiency against heterologous viral strains [12] and autologous HIV-1 typically evades contemporary antibodies through mutational escape [13]. This observation has led to uncertainty as to whether antibodies have a measurable impact upon HIV-1 infection dynamics [1]. Nevertheless, evidence that antibodies impact plasma viral load has been demonstrated in an HIV-1 infected individual in whom B-cells were depleted as treatment for lymphoplasmacytoid lymphoma [14]. In addition, broadly neutralising antibodies (bNABs), capable of binding diverse strains of HIV-1 have been identified in some individuals [15–19] and are now at the forefront of novel therapeutic applications (reviewed in [20]). Potential mechanisms for how antibodies impact HIV-1, *in vivo*, have been reviewed elsewhere [21] and include the following. 1) Antibodies could neutralize free virions, such that virus infectivity is impaired [22]. 2) Antibodies could bind infected cells and mediate antiviral activity, engaging Natural Killer cells for antibody-dependent cellular cytotoxicity or macrophages for phagocytosis and cellular destruction. 3) Immune complexes formed by antibodies bound to viral antigen could be taken up by dendritic cells, stimulating host adaptive immune responses. Identifying which of these mechanisms plays a measurable role in controlling HIV-1 remains to be elucidated.

A major obstacle in the development of a cure for HIV-1 is the presence of the latent reservoir [23–25]. This consists of a population of resting memory CD4+ T cells that can be maintained for many years in spite of combination antiretroviral therapy (cART) [26, 27] and lead to viral resurgence upon therapy cessation [28]. Progress towards a cure for HIV-1, requires us to understand more about how the latent reservoir is established and maintained. Until recently, the latent reservoir was thought to be established through a process in which cells are continuously deposited prior to cART; and maintained through a combination of proliferation [29, 30] and only sporadic reactivation of latent cells. In support of these assertions, the reservoir forms irrespective of whether cART is initiated early or late on during infection [31, 32]. Furthermore, both ultrasensitive measures of HIV-1 RNA in the plasma of patients on long term cART [8] and the fact that years of cART are insufficient for a cure, indicate that the reservoir decays very slowly indeed. Recent work however, has challenged these perceptions, [5, 6] revealing that the makeup of the latent reservoir typically matches viral strains that are circulating just prior to therapy. Only in a minority of individuals are the strains representative of those circulating throughout HIV-1 infection, pre-cART [6].

To explain these evolutionary findings, Abrahams et al [6] propose that cART alters the host environment in a way that allows the formation or stabilization of most of the long-lived latent HIV-1 reservoir. Furthermore, they describe how a mathematical model in which the turnover of latently infected cells is much faster prior to cART than during cART can explain how viral strains matching those circulating just prior to therapy are overrepresented in the reservoir. This new theory of reservoir dynamics has major implications for the prospects for an HIV-1 cure, pointing to new strategies that limit the formation of the reservoir around the time of therapy initiation. As such, there exists a need to interrogate whether other data support this theory and to further clarify it. For example, if the turnover of latently infected cells varies according to cART-driven alterations to the host environment, does this relate to a change in the rate at which latent cells are deposited, activated, lost (without activation) or proliferate?

In this study we analyse the viral and evolutionary dynamics of a unique patient who remained seronegative for 48 months following the onset of acute infection. By using a new mathematical framework at the interface of viral dynamic modelling and evolutionary modelling we address the following questions.

- How do the viral and evolutionary dynamics in this seronegative individual differ to those observed in seropositive individuals?
- What do these differences imply about

1. the role that antibodies play in controlling a typical (seropositive) HIV-1 infection.
2. the dynamics that underpin the formation or stabilization of the latent reservoir.

We highlight that although plasma viral load declined on cART, the rate of decline was considerably slower than in a typical seropositive individuals [6–8, 33] and that viremia remained detectable (850 copies/ml at its lowest) throughout the period prior to the onset of antibodies. By employing WGS analysis of longitudinal plasma samples, we further reveal a minimal increase in HIV-1 divergence during the period of measurable viremia, and absolutely no increase in divergence of nonsynonymous mutations during the first year of therapy. We show how a dynamical evolutionary within-host model of HIV-1 with the following two key features is the most parsimonious explanation for such minimal evolution in the face of measurable viremia. The first of these features is rapid activation of the latent reservoir prior to effective viral suppression. As such, our findings supports the theory presented recently by Abrahams et al [6] of rapid reservoir turnover prior to effective viral suppression. What is more, our findings elucidate this new theory by indicating that the rate of latent cell activation is the key component in this proposed rapid turnover. The second important feature of our model is that prior to the onset of antibodies in this patient, there are short chains of infection resulting from low levels of viral infectivity. This is consistent with the assertion that antibodies contribute to the suppression of viral infectivity in typical, seropositive individuals, perhaps through the direct neutralization of virions.

## Results

### Case summary and prior analysis

Our study focuses on a male, who had sex with men and who underwent routine HIV-1 screening every 6 months. Following infection (determined by both detection of viral RNA and DNA, and p24 antigen) he experienced the longest recorded time frame before seroconversion. Clinical details of this case have been previously reported [2]. Siemieniuk and colleagues reported that approximately 5 months prior to diagnosis, the patient was confirmed negative for HIV-1 through ELISA and retrospective PCR testing. Four months prior to diagnosis he experienced symptoms consistent with acute HIV-1. Over the next 4 months he was hospitalized twice with AIDS-defining illnesses. He was diagnosed HIV-1 positive through detection of proviral DNA by polymerase chain reaction (PCR), presence of HIV-1 p24 antigen and a plasma HIV-1 RNA viral load of 250,000 copies/ml. Viral load testing was conducted using the Roche Ultrasensitive Amplicor assay, version 1.5 and performed 39 times over a follow-up of 98 months. During this period, the patient was treated with combination antiretroviral therapy, including seven attempts to supress replication using recommended standard regimens. Viral loads declined approximately two logs between the start of cART and month 38 of cART and (atypically) remained at this detectable level until month 56. A weak HIV-1 specific antibody response was were first detected, during this interval, at month 48. By month 56, stronger antibody signals were detected. Overall, there is evidence that the patient had good adherence to cART. At month 62, however, the patient reported that – against medical advice – he was not taking cART. At that time, his viral loads were close to pre-cART levels. By month 67, the patient had started a new therapeutic regimen and viral loads had declined to undetectable levels, where they remained until the end of follow up. A timeline describing his infection is provided in Figure 1. The patient gave informed consent to have clinical data published in anonymity.

**Figure 1.**
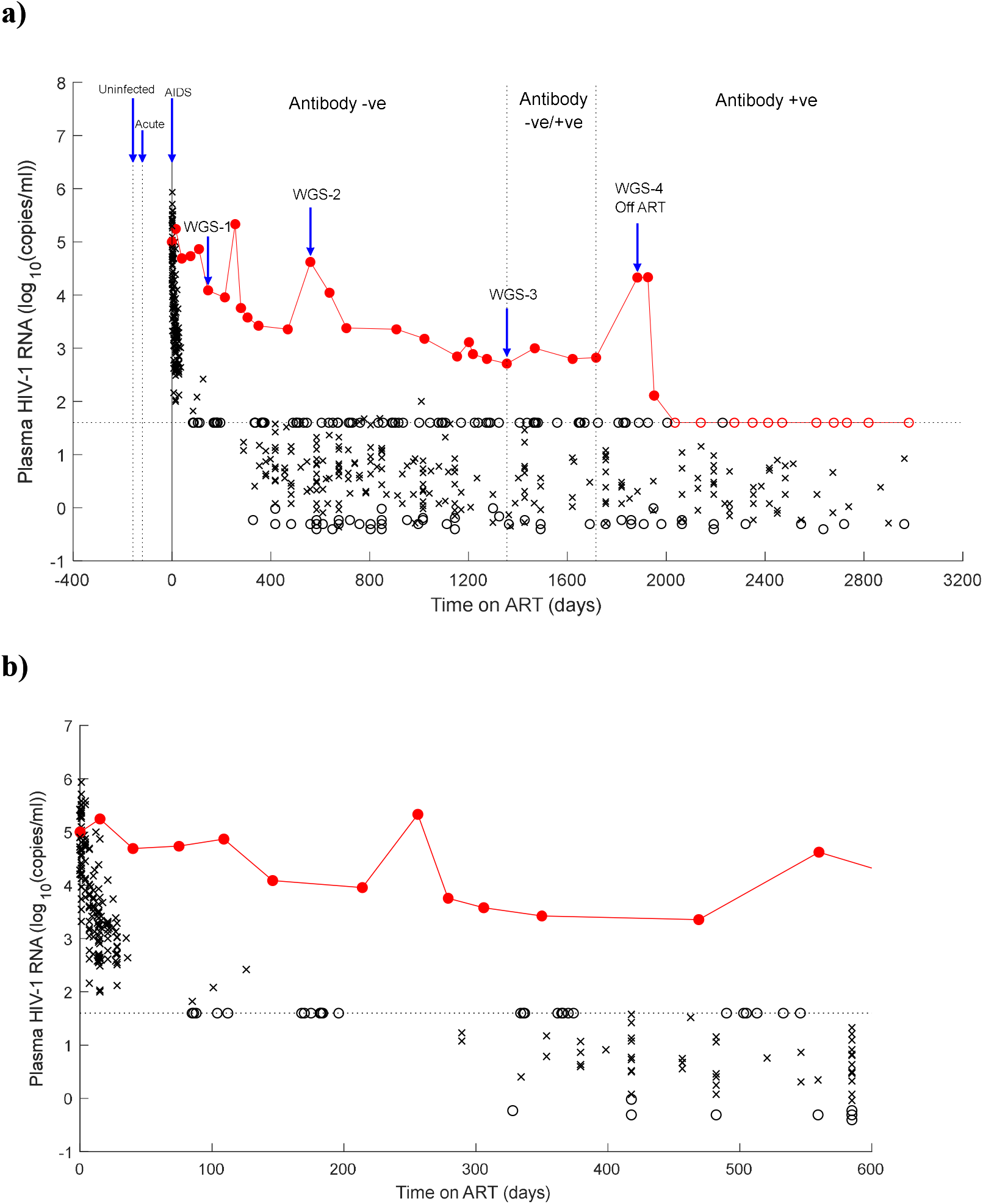
A comparison between the plasma viral load in the seronegative individual and in seropositive individuals undergoing cART. a) The comparison is made between 400 days prior to, and 3200 days post the onset of cART and is annotated to highlight key stages in the HIV-1 timeline. b) The comparison is made over the first 600 days of cART. Data from the seronegative patients are shown in red. Data from seropositive individuals have been extracted from two published studies [6, 8] and are shown in black. Filled red circles and black crosses represent detectable data. Unfilled black and red circles represent undetectable data and are plotted at the limit of detection. Blue arrows and vertical lines highlight key moments of the HIV timeline as follows: uninfected (day-158); infected with acute symptoms (day-120); day of diagnosis, AIDS symptoms and onset of cART (day 0); first WGS sample (day 146); second WGS sample (day 560); third WGS sample and weak anti-HIV-1 antibodies first detected (day 1355); strong anti-HIV-1 antibodies first detected (day x); fourth WGS sample (day 1883).

The cause of the prolonged period of seronegativity was investigated by looking at both host and viral factors [2]. Quantitative Ig and complement testing at the time of presentation did not show any evidence of immunodeficiency. The humoral deficit was shown to be specific to HIV: the patient mounted robust humoral responses to all challenge vaccines. The virus had similar gp120 antigenicity to HIV-positive control serum. Furthermore, genotypic resistance (Vircotype HIV-1, Janssen Diagnostics) testing of plasma RNA at 3, 10, 18, 44, and 61 months showed that the virus was pansensitive to non-nucleoside reverse transcriptase inhibitors (NNRTIs), nucleoside/nucleotide reverse transcriptase inhibitors (NRTIs), and protease inhibitors (PIs).

### Limited evolution despite years of measurable viremia

We aimed in this work to compare the viral and evolutionary dynamics of this long-term seronegative patient to those of ‘typical’ seropositive individuals. Figure 1 compares the viral dynamics of this patient to those of 49 seropositive individuals undergoing cART [6, 8, 33]. It highlights that in seropositive individuals, viral loads decline with a mean half-life of 1.5 days during the first week of cART [28, 34–36] and a mean half-life of 28 days during the next few weeks. [8] Furthermore, they are typically undetectable (<30 copies/ml) using a standard assay after 200 days of cART, and stabilize to a mean value of 2 copies/ml after 320 weeks of cART [8]. By contrast, in the seronegative patient, although viral loads markedly declined on therapy in the absence of antibodies, both the gradient and the longer-term magnitude of the decline was much more modest. Only after antibodies had appeared, did viremia further reduce, by orders of magnitude, to undetectable levels.

To investigate virus evolution, we performed whole genome next generation sequencing of longitudinal plasma HIV-1 RNA samples from the seronegative patient. Samples were collected at four times when viremia was measurable: approximately 4, 18, 46 and 62 months post diagnosis. Paired-end sequencing was performed on the MiSeq Illumina platform. To infer the patient’s whole genome consensus sequence at each time point, short reads were assembled de-novo. Trimmed reads were mapped to the patient’s *de novo* assembled HIV-1 genome and deduplicated. The mean length of the merged reads (across all time points) was 135 nucleotide sites (Table S1). We focussed our evolutionary analysis on four regions of the HIV-1 genome: gag, pol, env and nef. Mean coverage across these regions was 647, 879, 66 and 1565 reads at 4, 18, 46 and 62 months post diagnosis, respectively (Table S1). Due to the low coverage at 46 months post diagnosis, data from this sample time were excluded from our analysis, except where we explicitly state otherwise. At each of the three remaining sample times, and for each of the four gene regions, we inferred the mean synonymous and nonsynonymous divergence away from the consensus sequence at 4 months post diagnosis, as well as the mean diversity (see Methods). Because all our samples were taken at times when viremia was measurable, we compared these evolutionary metrics to those observed in a group of nine untreated seropositive HIV-1 infected individuals who have been studied in detail elsewhere [4].

Figure 2 reveals that diversity in the seronegative patient at the first sample time (day 146 of cART) was either above (gag and pol and; Fig 2a and 2b) or close to (env and nef; Fig 2c, 2d) the high end of the distribution of diversity observed in the seropositive individuals. A plausible explanation for this is that the seronegative patient was infected with more than one strain. By contrast, both synonymous (Fig 3a-d) and nonsynomymous (Fig 3e-h) divergence at the first sample time fell within the distribution exhibited by seropositive individuals for all four gene regions, with one exception (nonsynonymous divergence in pol; Fig 3f). Strikingly, over the next 14 months, measures of evolution were minimal. Synonymous divergence either declined (gag and pol; Fig 3a-b and Fig 4a) or remained approximately constant (env and nef; Fig 3c-d and Fig 4a), in contrast to the increase in divergence typically exhibited by the seropositive individuals over the same period. Increases in diversity (in gag, env and nef; Fig 2) and nonsynonymous divergence (in all genes; Fig 3e-h) were observed, but the rate that synonymous divergence accrued over this period was small compared to that observed in the untreated seropositive individuals (Fig 4b). To further investigate evidence for evolution, synonymous and nonsynonymous ‘sweeps’ were also identified (Fig S1, Fig S2 and Table S2). These were defined when substitution at a particular site away from the consensus nucleotide at the first sample time, was greater than 50% at any later sample time (see Methods). Five such sites, (all nonsynonymous) were identified by 18 months post diagnosis. Arguably, the minimal degree of evolution observed during this 14 month period of persistently measurable viremia, is unexpected. This is because measurable viral loads are typically attributed to ongoing cycles of replication and are associated with positive measures of evolution.

**Figure 2.**
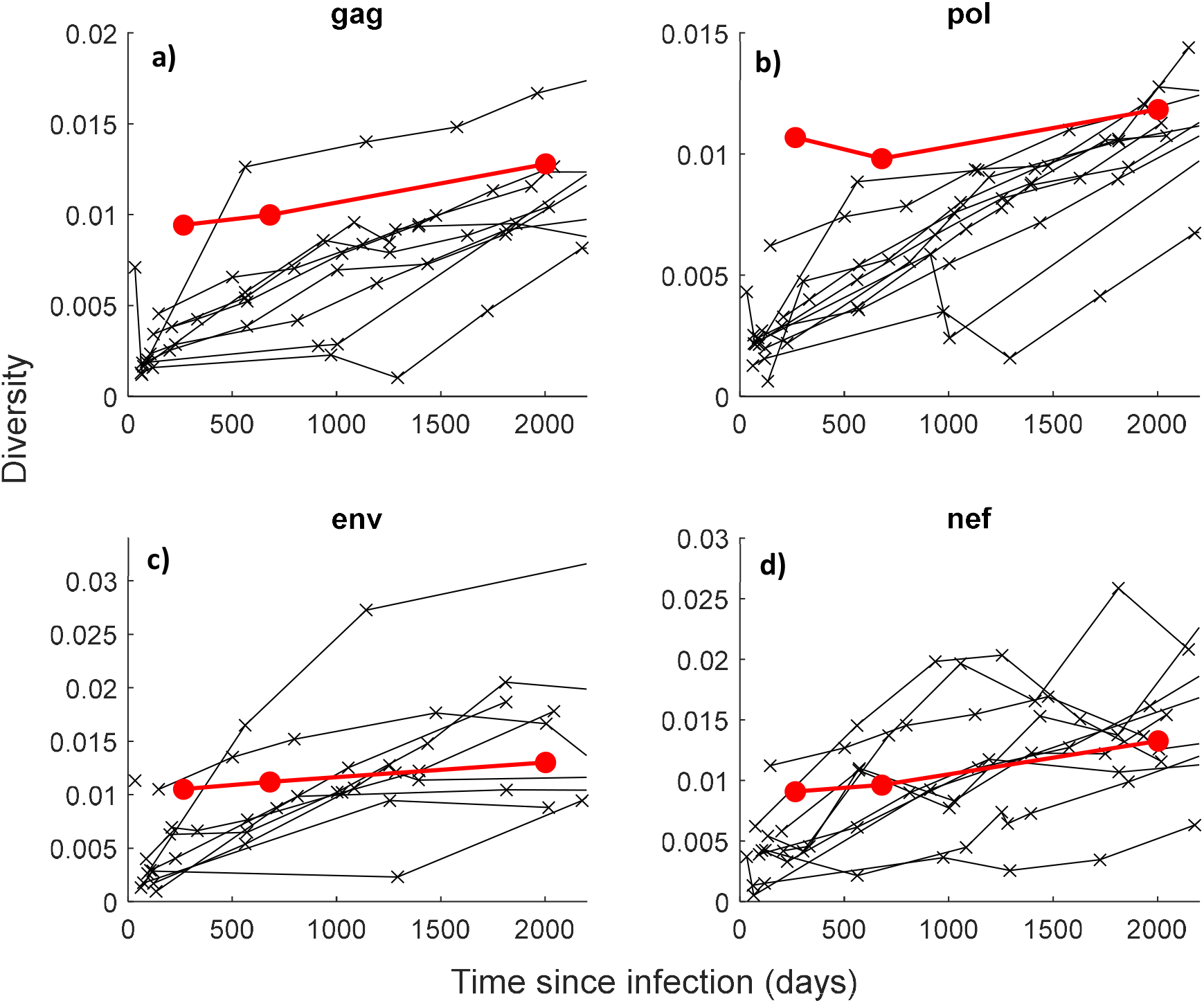
A comparison between diversity in the seronegative patient and untreated seropositive individuals. Mean diversity over the first 2000 since infection is compared between the seronegative individual (red markers) and nine untreated seropositive individuals (black markers). These data and are stratified according to gene: gag (a and e), pol (b and f), env (c and g) and nef (d and h).

**Figure 3.**
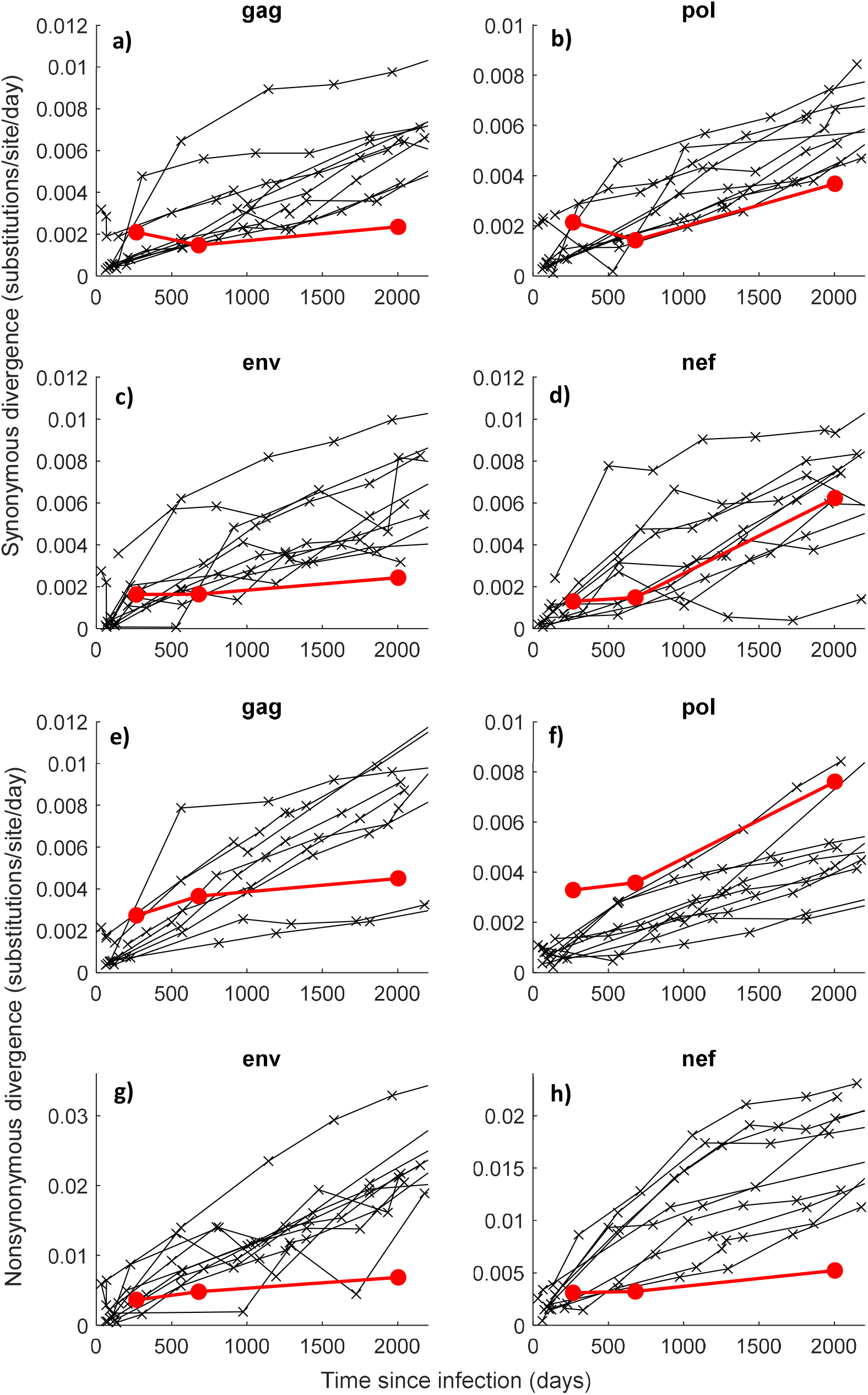
A comparison between divergence in the seronegative patient and nine untreated seropositive individuals. Synonymous (a-d) and nonsynonymous divergence (e-f) are compared between the seronegative individual (red markers) and nine untreated seropositive individuals (black markers)[6]. These data are shown for the first 2000 days post infection and are stratified according to gene: gag (a and e), pol (b and f), env (c and g) and nef (d and h).

**Figure 4.**
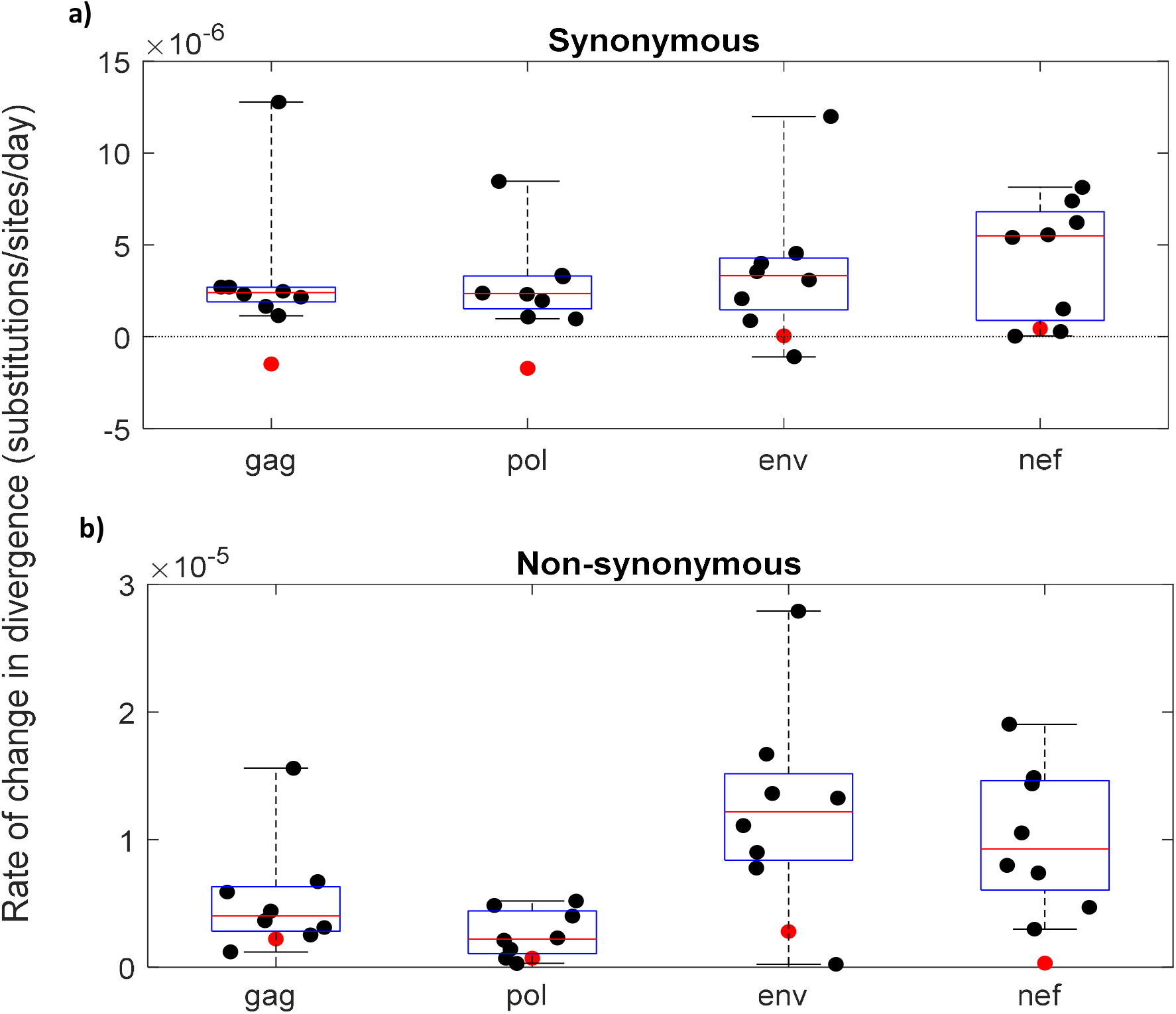
The rate of change in divergence between days 266 and day 580 post infection in the seronegative patient is compared to the change in divergence over a similar timeframe in the untreated seropositive individuals. The rate of change in synonymous (a) and nonsynonymous divergence (b) from day 266 to day 580 post infection in the seronegative individual (red markers) is compared to the rate of change in divergence over a similar timeframe in nine untreated seropositive individuals (black markers)[4]. Because the seropositive individuals are not sampled at exactly same times as the seronegative individual, the change in the divergence in seropositive individuals is estimated using linear regression over the shortest time period spanning the period day 266 to day 580 post infection. The data are stratified according to gene. The boxplots show the median (red lines), interquartile range (blue box) and full range (black lines) of the rate of change in divergence amongst the seropositive individuals.

Between month 18 and month 62 post diagnosis, there was a measurable degree of evolution, characterised by increasing diversity (Fig 2), synonymous divergence and nonsynonymous divergence across all genes (Fig 3). However, synonymous divergence in gag and env (Fig S3a); nonsynonymous divergence in gag, env and nef (Fig S3b); and diversity in env (Fig S4) all increased at rates that were very low compared with those observed in untreated individuals. In total, sweeps at 37 nucleotide sites (25 nonsynonymous and 12 synonymous) were identified by month 62 post diagnosis (Fig S1, Fig S2 and Table S1). Furthermore, only two of these sweeps (both nonsynonymous in env) exhibited a measurable substitution prevalence at month 62 post diagnosis. Arguably, if the measurable viral loads observed in this patient were the results of continuous cycles of ongoing replication, such sweeps – which indicate evolutionary progression of the virus – would be more prevalent.

### A new within-host mathematical model of HIV-1 infection

We aimed to investigate what the atypical viral and evolutionary dynamics observed in the seronegative patient tell us about HIV-1 biology. Specifically, we focussed on what these dynamics suggest about: 1) the dynamics that underpin the formation of the latent reservoir. and 2) the role that antibodies play in controlling typical (seropositive) HIV-1 infection. To achieve this goal, we developed a novel within-host model of HIV-1 at the interface of viral dynamics and evolutionary modelling that can simultaneously explain the dynamics observed in the seronegative individual and in typical seropositive individuals.

Our model provides a new platform for investigating the interplay between HIV-1 viral and evolutionary dynamics and is detailed in the Methods and Figure 5. The backbone of the model is a within-host description of HIV-1 infection that tracks the infection of cells by free virus [7, 30, 37, 38]. In brief, there are two classes of target (CD4+) T cells, which become infected with HIV-1 as a result of interaction with free virus. Most target cells facilitate rapid integration; the majority of new infections amongst this class result in productively infected cells with a short (~1.5 days) half-life and the remainder result in latently infected cells. A smaller class of target cells facilitate slower integration; upon infection these cells initially enter a pre-integration stage with a comparatively long half-life (~28 days). Both latently infected cells and pre-integration infected cells can transition to a state of productive infection, associated with a short half-life. Antiretroviral therapy is assumed to contribute to reducing the rate that new cells are infected.

**Figure 5.**
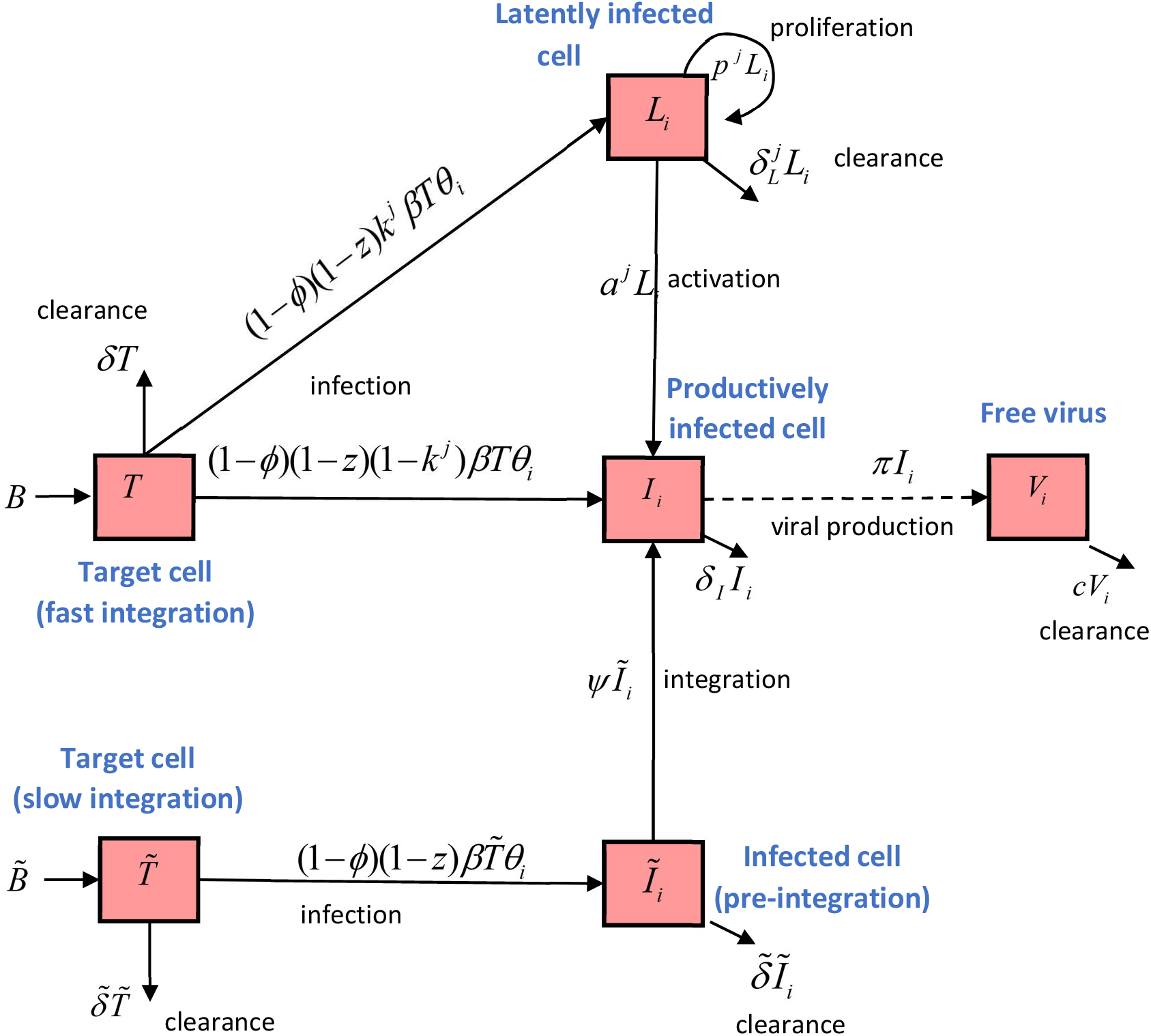
A within-host mathematical model of the spread of HIV-1 and strain evolution. A description of the model is provided in Methods.

In addition to a standard backbone, our model has three key components. First, it accounts for a contribution from anti-HIV-1 antibodies in reducing viral infectivity. Second, it allows for variability, according to whether viral replication is effectively suppressed, in parameters that govern the dynamics of latently infected cells. These include a parameter that determines the rate that latent cells are generated, and rate parameters of latent cell clearance, activation (which results in short-lived, virus-producing infected cells) and proliferation (which maintains the viral strain). Third, it incorporates the evolution of viral strain divergence away from a founding strain. During each infection cycle, neutral substitutions accrue along the genome at a rate proportional to the number of substitution-free sites.

### Model implications for evaluating reservoir formation and maintenance

We first used our model to investigate the dynamics that underpin the formation of the latent reservoir. Our model is a flexible extension of classical within-host dynamical models of HIV-1 that have been shown to explain key aspects of the viral dynamics of (typical) seropositive individuals [7, 30, 37, 38]. Central to these classical models is that the differing decay rates of free virus and different classes of infected cells provides an explanation for why viremia appears to decay in multiple phases following the onset of cART. When our model is parametrized to represent infection in a typical seropositive individual – through an assumption that antibodies are present and reduce viral infectivity – our model is similarly able to reproduce these dynamics (Text S1). Specifically, the inclusion of free virus, as well as short-lived, long-lived (pre-integration) and latently infected cells predicts that virus decays in four phases, with the last phase decaying with a very long half-life, representing the decay of the latent reservoir.

As our model tracks how neutral substitutions accrue along the genome over time, it is additionally able to recreate a key evolutionary pattern that has been observed in untreated seropositive individuals; namely a gradual increase in the mean nonsynonymous divergence over time (Text S1). In addition, the model can predict how nonsynonymous divergence changes once cART is started in seropositive individuals. Traditionally, models of HIV-1 dynamics assume that the turnover of latently infected cells is slow, both before, and during, the onset of cART. When our model is parametrized in this manner, it predicts that the mean divergence of virus in plasma is significantly reduced once cART has started (Text S1). This agrees with the long-perceived wisdom that the reservoir contains an archive of viral strains that, on average, are older than the strains that were circulating in plasma around the time that cART begins. In such a scenario, the small levels of virus that can be found in plasma during cART derive from reactivation of latent viral strains and would be expected to be less diverged than virus (derived from active replication) present in the plasma at the time of cART onset (Text S1).

Recent findings, however, are leading the field to revaluate the theory that the reservoir contains an archive of viral strains. Two studies have now revealed that viral strains that come from the reservoir after several years of cART are, in most people, genetically most similar to those circulating *close* to the time therapy started [5, 6]. In one of these studies [6], a model in which the ‘turnover’ of the latent reservoir is faster prior to the start of therapy than during therapy, was provided to explain these seminal evolutionary findings. However, the model was not explicitly described in that report and clarification and further examination of the proposed mechanisms involved are therefore necessary. Exploring the mechanisms that relate to the formation and maintenance of the latent reservoir may provide direction for future cure initiatives.

In Text S1 we describe how we have examined these mechanisms explicitly by fitting our model to viral and evolutionary dynamics that are typical of seropositive individuals. We focus on whether variation, according to whether virus replication is effectively suppressed (governed by cART status), in the rate that latently infected cells are produced, cleared, activated or proliferate, can explain these new evolutionary findings. We highlight that (nonexclusive) potential explanations are that the rate that latently infected cells are produced or proliferate is slower prior to cART compared to during cART, or that the rate that latently infected cells are activated or cleared is faster prior to cART compared to during cART. In light of these model results, alternative data sources are therefore necessary to further explore for which of these rates, there exist evidence of variability across infection, according to factors that include treatment status.

### Our study supports the rapid activation of latently infected cells prior to effective viral suppression and a role for antibodies in reducing viral infectivity

By fitting our model to the viral and evolutionary dynamics of the seronegative individual, we investigated the role that antibodies play in controlling a typical (seropositive) HIV-1 infection and further explored the dynamics that underpin the formation of the latent reservoir. Specifically, we investigated the source of variation – between treated and untreated infection that governs the dynamics of latently infected cells in the context of typical seropositive individuals.

Several features of the viral and evolutionary dynamics of the seronegative individual are noteworthy and, when explored in the context of our model, inform these areas of study. The first feature is that in the absence of an antibody response, plasma RNA viral load reduced by orders of magnitude following the onset of drug therapy, but remained detectable (and above 500 RNA copies/ml), even after 46 months of cART. However, following the emergence of a robust antibody response, plasma RNA viral load further reduced by orders of magnitude until it was undetectable. Through exploration of the model, we reveal that the most parsimonious explanation for these data is that antibodies typically contribute to reducing viral infectivity. As such, in their absence, at least some degree of viral infectivity – and new infections of cells – persists.

The second key feature of the data is that there was almost no increase in viral strain divergence between months 4 and 18 post cART initiation and only a limited increase in strain divergence between months 18 and 62 post cART initiation, despite measurable viremia across both these of these intervals. Of note, some fluctuations in viral load were observed over these time intervals, with the most dramatic increase in viral load measured at month 62 when the patient reported that (against medical advice) he had stopped taking cART. This second feature can be explained by two alternative hypothesis. One hypothesis is that ongoing cycles of replication were principally responsible for the measurable levels of viremia, but substitution rates were atypically low in this patient. In this hypothesis, viral infectivity is assumed to be sufficiently high in the absence of antibodies (but presence of cART), that the within-host reproductive number of HIV-1 is consistently above unity. A second hypothesis is that, the within-host reproductive number was, instead, mostly below unity, meaning that most virus does not result from ‘active’, continuous cycles of replication. Instead, central to the virus observed in the plasma is the reactivation of latently infected cells. Some of the virus will have resulted directly from reactivation and some will have resulted from chains of cell infections that start with reactivated virus and terminate after a short number of rounds. Explaining the non-zero (albeit minimal) measures of evolution, some may even have resulted from longer chains of infections that occurred during short periods where the within-host reproductive number fluctuated above unity (e.g. due to varying cART adherence). Fundamentally, though, the limited degree of evolution relates to the assumption that the strains that were sampled during therapy are closely linked to ones that were archived prior to therapy.

Although we cannot rule out the first of these hypotheses (ongoing replication cycles), it relies on the seronegative individual exhibiting atypically low substitution rates. By fitting our model to the viral dynamics and synonymous divergence data from the seronegative patient in a manner that is also consistent with data from typical seropositive individuals (see Methods and Text S1 for details), we show that that the second of these hypotheses is a more parsimonious explanation of the data (see Figure 6 for the least squares optimal fit). We note that this second hypothesis is consistent with an assumption that the level of virus that derives directly from activation of latently infected cells in the cART-treated seronegative individual (prior to the onset of anti HIV-1 antibodies) is larger than is typical in cART-treated seropositive individuals – where viremia is normally undetectable. Thus, it supports the findings of Abrahams et al [6], that the turnover of the latent reservoir (see equation 1) is more rapid prior to, than post, effective viral suppression. What is more, it shines a new light on their findings, by indicating that the rate of latent cell reactivation may contribute to this source of variation in ‘turnover’.

**Figure 6.**
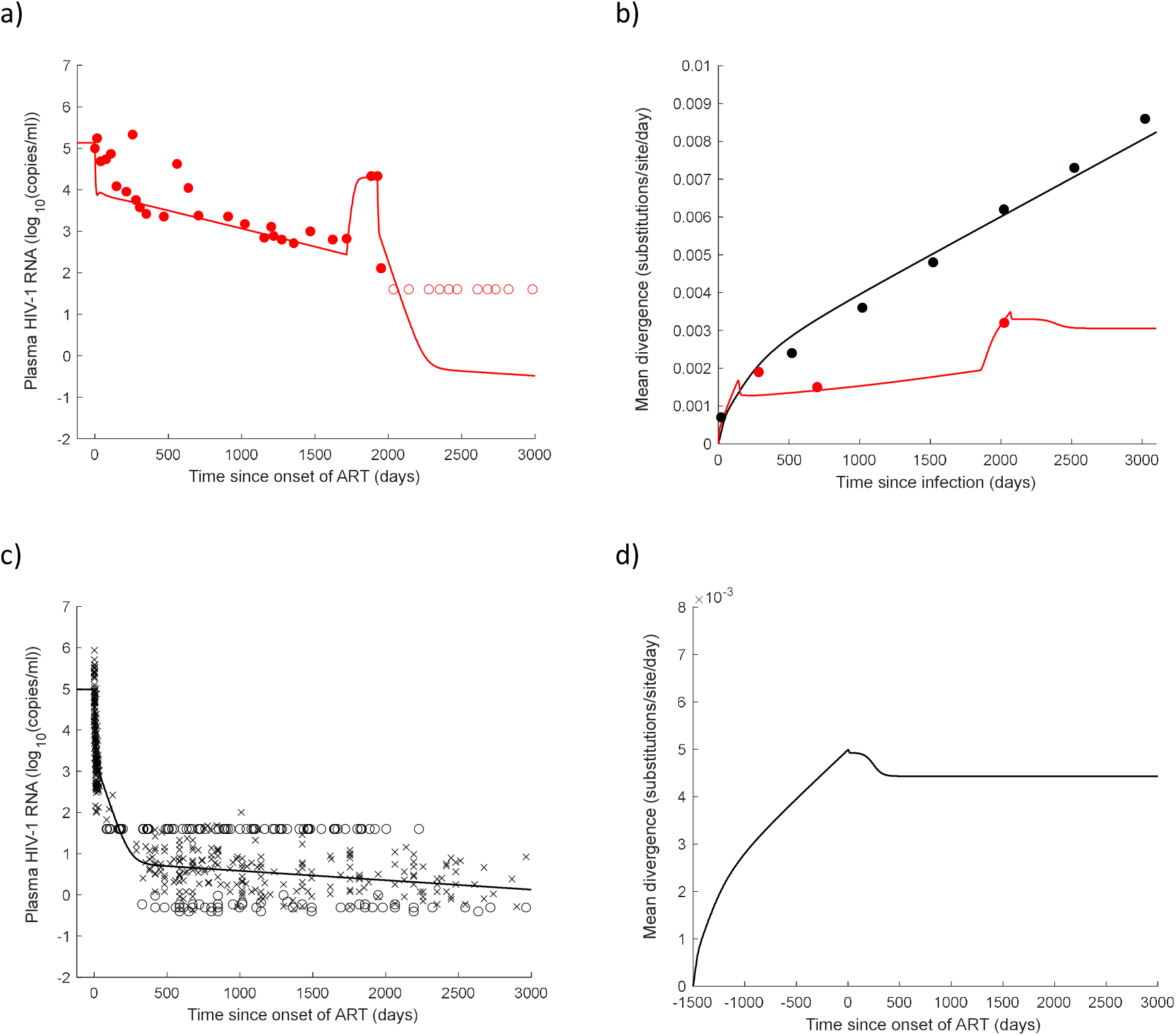
A model in which 1) antibodies contribute to suppressing viral infectivity and 2) the activation rate of latent cells is faster before, compared to after, viral replication is effectively supressed, can simultaneously explain viral and evolutionary dynamics observed in the seronegative individual and typical seropositive individuals. Optimal model parameters are inferred by simultaneously fitting the model to data from the seronegative individual and to seropositive individuals under different cART and antibody scenarios. In a) optimal model predictions of plasma viral loads are compared to those observed in the seronegative individual. In b) optimal model predictions of mean divergence are compared to mean synonymous divergence (averaged across the four genes: gag, pol, env and nef) observed in the seronegative individual (red) and in untreated seropositive individuals (black) [4]. In c) the optimal model predictions of plasma viral loads are compared to those observed in treated seropositive individuals [6, 8]. In d) the optimal model predictions of mean divergence following the onset of cART in seropositive individuals are shown. This reveals that divergence on cART is close to the mean divergence of strains present in the plasma close to the time that cART starts in accordance with recent findings [6].

## Discussion

In this study we developed a new within-host viral dynamics and evolutionary model of HIV-1. With this model, we first highlighted how a recently supported hypothesis [6] that viral strains in the reservoir turnover more rapidly prior to, than post, viral suppression is consistent with both the viral and evolutionary dynamics of typical seropositive individuals. Furthermore, we showed how an increase in the rate of production or proliferation, or a reduction in the rate of clearance or activation of latently infected cells are all potential explanations for such a shift and further pinpointing *which* of these rates change will be critical to novel HIV-1 cure initiatives.

The principal focus of this study, however, was to analyse the viral and evolutionary dynamics of HIV-1, sampled from the plasma of an individual who failed to seroconvert until the month 48 of cART. Whole genome sequencing revealed minimal measures of evolution, despite years of measurable viremia during cART in this unique individual. Using our mathematical model we demonstrated how our data are consistent with a theory that once antibodies are present in HIV-1 infected individuals, they play a measurable role in reducing new infections of cells, perhaps through binding to virus in such a way that they block virus-target cell binding. This agrees with, and provides a potential explanation for, a previous finding that antibodies can impact HIV-1 plasma viral load [14]. By improving our understanding of the role that antibodies play in controlling natural HIV-1 infection, this study adds support for antibody-based vaccine and therapy development [21].

We note that an alternative model formulation in which antibodies reduce viral production could equally well reproduce a key feature of the viral dynamics in the seronegative patient – the orders of magnitude reduction in viral load following the appearance of antibodies (Text S1). However, the biological mechanism behind such a potential reduction in viral production is less clear, and thus we do not promote such a theory. We also acknowledge that our model does not include the potential impact that antibodies could have on viral or cell clearance. Although our model would remain consistent with the key features of the data if the direct impact of antibodies on clearance rates were additionally included, these mechanisms alone are insufficient to explain the orders of magnitude reduction in viral load following the appearance of antibodies, which the assumption of a reduction in viral infectivity does predict (Text S1). For this reason we do not focus on the role of antibodies on viral and cell clearance rates.

By fitting our model to the data from the seronegative patient, we further predict that in the absence of antibodies (but presence of cART), rapid activation of latently infected cells and some short chains of infections that ultimately terminate, yield measurable levels of virus in blood, but no substantial increase in divergence over time. Our study supports a hypothesis that a reduction in resting CD4+ T-cell activation contributes to the reduction in the turnover of viral strains in the latent reservoir that has been demonstrated in seropositive individuals in response to effective viral suppression [6]. A possible explanation for this is that the immune activation and dysfunction associated with viremia and the lack of IL-7/IL-7R signalling through CD127 precludes effective memory maintenance [39]. As such, these cells activate, on average, after a short period of time, i.e. they are only short-lived memory CD4+ T-cells. By contrast, in the context of effectively supressed viral replication, the reduction in immune activation and associated restoration of IL-7/IL-7R signalling through CD127 provides the environment for memory CD4+ T-cells to be maintained as long-lived cells [39]. Thus, whilst some sporadic activation of these memory CD4+ T cells may occur once viremia is supressed, on average their rate of activation is much slower. It is an exciting prospect that novel therapeutic strategies that combine cART initiation with inhibition of IL-7/IL-7R signalling to limit CD4+ T cell memory maintenance could limit the HIV-1 reservoir and provide a path to a functional cure [40].

We stress that our findings are based upon data obtained from only a single individual and therefore require further confirmation. The individual studied here is unique in the duration of identified seronegativity; however, other individuals with shorter periods of seronegativity have been identified and offer a potential avenue for study. What is more, our study highlights the value of identifying new seronegative individuals and the application of modelling to simultaneously examine their viral and evolutionary dynamics.

We emphasize that our modelling framework has broader potential than just the application presented here. Natural extensions to the model include the tracking of adaptive (synonymous) substitutions that can lead to selective sweeps and the addition of other biological compartments under investigation, such as drug sanctuaries and cellular subspecies. Here, the evolutionary output that we have focussed on is the mean divergence, however the model can also predict the distribution of the divergence of viral strains, which could help test hypotheses, such as whether ongoing cycles of replication persist in drug sanctuaries during apparently suppressive cART.

The prospect that biological processes pivotal to the establishment of the reservoir are only operating at their peak once cART has started, implies an enormous opportunity for cure research to shift focus from eliminating the reservoir, to preventing its formation. Mathematical models at the interface of viral dynamics and evolutionary modelling have an important role to play in the critical examination of future trials data to further advance the development of a cure.

## Methods

### Sequencing

Viral RNA was extracted from the patients’ serum using a QIAamp Viral RNA Mini Kit (Qiagen, Hilden, Germany). Paired-end sequencing was performed by the University College London Pathogen Genomics Unit: (https://www.ucl.ac.uk/infection-immunity/pathogen-genomics-unit) on the MiSeq Illumina platform using MiSeq micro 300 run kit. The library preparation was performed using the Agilent RNA SureSelectXT kit with HIV specific baits.

### Bioinformatics

Raw sequence reads were pre-processed using Kraken v1 [41] to identify contaminant (human and bacterial) sequences, using a custom database containing all bacterial, viral and archaeal genomes from RefSeq and the human genome. Sequences identified as viral or unclassified by Kraken were used for assembly. To infer the patient’s HIV-1 consensus sequence at the first timepoint, reads were assembled with shiver [42], after using SPAdes v 3.10 [43] to generate the contigs. The resulting assembled HIV-1 genome sequence was curated manually in AliView, and used as the mapping reference for all reads from subsequent timepoints. For each timepoint, reads were quality-trimmed and stripped of adaptors with bbduk (sourceforge.net/projects/bbmap/) (parameters: ktrim=r k=23 mink=11 hdist=1 minlen=50 qtrim=rl trimq=30 tpe tbo), before being mapped to the patient’s custom reference sequence using bowtie2 (http://bowtie-bio.sourceforge.net/bowtie2). Mapped reads were deduplicated using MarkDuplicates from the Picard software package (http://broadinstitute.github.io/picard). BAM files were converted to SAM using samtools [44], and used to generate fasta alignment files with a custom-made java program.

### Synonymous and nonsynonymous divergence

Synonymous and nonsynonymous divergence at each sample time and for each of the four gene regions was calculated. Codons with a read depth of at least 50 at each of the three nucleotide sites at both that sample time point and the first sample time (4 months post diagnosis) were first identified. At each such codon, the amino acid corresponding to the consensus sequence at the first sample time was inferred. At each sample time, frequencies of each of the alleles A, C, G and T alleles at each site were then calculated and used to infer the frequencies of all possible triplets – and corresponding amino acids – at each codon. These frequencies were then used to infer the absolute prevalence of synonymous and nonsynonymous polymorphisms at each codon. By finding the mean prevalence across all data-sufficient codons in the gene and dividing by three, the mean synonymous and nonsynonymous divergence (per nucleotide site) was inferred.

### Nucleotide diversity

To estimate nucleotide diversity, sites with a read depth of at least 50 were first identified. At each such locus, l, the proportion of pairwise differences *D_l_* between alleles was then inferred as follows. Let *N_l_* be the read depth at locus l and *A_l_*, *C_l_*, *G_l_* and *T_l_* be the number of copies of each of the four alleles at locus *l*.

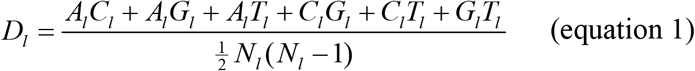

The diversity statistic, π, was then calculated by finding the mean proportion of pairwise differences across all nucleotide sites in the gene [45].

### Identifying sweeps

To identify sweeps, nucleotide sites with a read depth of at least 50 at the first sample time (4 months post diagnosis) and at least one other sample time were first identified. At each such locus and sample time with sufficient read depth, the prevalence of polymorphisms away from the consensus allele at the first sample that correspond to synonymous and nonsynonymous changes are calculated. ‘Sweeps’ were identified at loci where a polymorphism prevalence greater than 50% was achieved at least one sample time. Because we restricted our analysis to sites with sufficient read depth, we did not explicitly compare the number of sweeps identified in the seronegative patient to the number seen in the Neher dataset, where coverage was higher.

### A mathematical model of within-host HIV-1 viral dynamics and strain evolution

The variable *t* denotes the time (in days) and the index *j* denotes whether virus replication is effectively suppressed (supressed: j = 1 and not supressed: j = 0). There are two classes of uninfected target cells. Uninfected target cells that facilitate rapid and slow integration (*T*(*t*)) are assumed to enter the system at rate *B* days^-1^ and die at a per cell rate of *δ* days^-1^. Uninfected target cells that facilitate slow integration 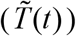 are assumed to enter the system at rate 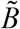 days^-1^ and die at a per cell rate of 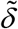 days^-1^. The index *i*, denotes the class of viral strains that have substitutions away from the founding strain at *i* of the n sites in the viral genome. In the absence of either treatment or an antibody response target cells become infected with an i-substitution strain at a per cell rate that is equal to the product of the transmission parameter *β*, and the variable *θ_i_*(*t*), which is proportional to the fraction of new infections at time *t* with an *i*-substitution strain. In the presence of antiretroviral therapy, this rate is reduced by a proportion 1 − *z*, where *z* is defined as the effective drug concentration. In the presence of an antibody response, the rate is further reduced by a proportion 1 − *ϕ*, where *ϕ* is defined as the effectiveness of antibodies in reducing viral infectivity. A fraction *k^j^* of all newly infected cells that facilitate rapid integration, become latently infected (*L_i_*(*t*)). These cells do not produce free virus and are cleared at per cell rate of 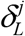 days^-1^, proliferate at a per cell rate of *p^j^* days^-1^ and are activated into a state of productive infection at a per cell rate of *a^j^* days^-1^. We assume that cells that are created as a result of proliferation retain the viral strain of the proliferating cell. The model has the flexibility to allow these parameters, governing reservoir formation and maintenance (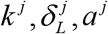 and *p^j^*), to be dependent upon whether virus replication is effectively supressed or not (indexed by *j*). The remaining fraction (1 − *k^j^*) of all newly infected cells that facilitate rapid integration become productively infected (*I_i_*(*t*)), whereby they produce virus with an *i*-substitution strain (*V_i_*(*t*)) at a per cell rate of *π* days^-1^. These cells are cleared at a per cell rate of *δ_I_* days^-1^ and free virus is cleared at a per virion rate of *c* days^-1^. All newly infected cells that facilitate slow integration, first enter an unintegrated infected state, from which they are cleared at a per cell rate of 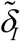 days^-1^ and undergo integration, facilitating productive infection, at a per cell rate of *ψ* days^-1^. A fraction, *μ*(*n* − *i*) of new infections that are caused by the interaction of target cells with infectious free virus with *i*-substitutions have a single additional substitution. Here, *μ* is the per site probability of substitution upon cell infection and *n* − *i* is the number of sites at which substitutions are still possible. The remaining fraction of new infections, 1 − *μ*(*n* − *i*), have no additional mutations. As the rate of substitutions at two or more sites per cell infection is low for HIV-1 (order *μ*^2^) we assume that double mutations do not occur. We further assume that ‘backwards’ substitutions (substitutions towards the founding strain) do not occur and that substitutions have no impact on viral fitness. These reactions are described by a set of coupled ordinary differential equations (equations 2–7) and an expression for the variable *θ_i_*(*t*) (equation 8).

### Model fitting

By employing a least squares approach (using the Matlab function lsqcurvefit), we have fitted the mathematical model to longitudinal data relating to plasma HIV-1 RNA and synonymous divergence from the seronegative individual. It was simultaneously constrained to ensure that when the antibody response (governed by parameter *ϕ*) is present in the model, the model is consistent with viral and evolutionary dynamics data from typical seropositive individuals. To fit the model to plasma HIV-1 RNA viral loads, the number of copies/ml were scaled by 7500 to account for the assumptions that two RNA copies exist in each virus particle and that there is a total 15 litres of extracellular fluid in the body. The following parameters were fixed to published values: 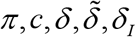 and *ψ*– this did not impact our findings. The total number of sites, n, was chosen to be sufficiently large that saturation of mutations within the genetic region does not measurable impact the accrual of genetic divergence. The substitution rate parameter, *μ*, was estimated by fitting a model that characterized an untreated, seropositive individual to a constant plasma HIV-1 viral load of 5 log_10_ copies/ml and to longitudinal data of the mean (averaged across gag, pol, env and nef) synonymous divergence inferred from untreated seropositive individuals (Fig 6). The target cell production rate parameter, *B*, was estimated by fitting a model that characterized an untreated seronegative individual to the plasma viral load of the seronegative individual at diagnosis. The transmission parameter, *β*, was then inferred by fixing the basic reproductive number (R_0_) to a value supported by data using the expression *β =*(*R_0_δδ_I_c*)/(*Bπ*), which is a good approximation to the basic reproductive number for this system. All remaining model parameters were estimated by simultaneously fitting the model to the plasma HIV-1 RNA and synonymous divergence data from the seronegative individual under the following constraints. 1) Parameters that govern the rates of latent cell production, proliferation and clearance are assumed to be independent of whether viral replication is effectively supressed: *k*^0^ = *k*^1^, *p*^0^ = *p*^1^ and 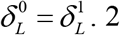) When the model is parametrised to represent a treated seropositive individual, plasma viral load is fitted to equal 1.5 copies/ml after 360 weeks of cART [8] and the mean divergence of virus in plasma during cART is fitted to equal the mean divergence of virus in plasma two thirds of a year prior to the start of cART, in agreement with data indicating that most viral sequences from the reservoir are most closely related to sequences present in the plasma within the year preceding cART. [6]. 3) The long-term decay rate of virus in plasma when viral replication is effectively supressed 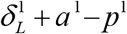 is fixed to a value supported by data. Refer to Table 2 for parameter values and references.

**Table 1.** Model variables

| Variable | Description | Start conditions<br>( $t = 0$ ) |
| --- | --- | --- |
| $t$ | Time (days) | |
| $j$ | Index identifying whether viral replication is effectively suppressed ( $j = 0$ not suppressed, $j = 1$ suppressed) | |
| $i$ | Number of mutated sites away from the index strain, $I_0$ | |
| $\theta_i(t)$ | Fraction of new infections with a strain with $i$ mutated sites at time $t$ | |
| $T(t)$ | Number of target cells at time $t$ that facilitate rapid integration | $B/\delta$ |
| $\tilde{T}(t)$ | Number of target cells at time $t$ that facilitate slow integration | $\tilde{B}/\tilde{\delta}$ |
| $I_i(t)$ | Number of cells that are productively infected with a strain with $i$ mutated sites at time $t$ | 1 $i = 0$<br>0 $i = 1 : n$ |
| $\tilde{I}_i(t)$ | Number of slow-to-integrate cells that are in a pre-integration infected state with a strain with $i$ mutated sites at time $t$ | 0 $i = 0 : n$ |
| $L_i(t)$ | Number of cells that are latently infected with a strain with $i$ mutated sites at time $t$ | 0 $i = 0 : n$ |
| $V_i(t)$ | Number of free virions with a strain with $i$ mutated sites at time $t$ | 0 $i = 0 : n$ |

**Table 2.** Model parameters

| Parameter | Description | Value used | Reference |
| --- | --- | --- | --- |
| $B$ | Rate of production of target cells that facilitate rapid integration | 3525000 | Fitted |
| $\tilde{B}$ | Rate of production of at which target cells that facilitate slow integration | 35250 | Inferred from [8] |
| $\delta$ | Per cell clearance rate of target cells that facilitate rapid integration | 0.15 | <sup>a</sup> Inferred from [28, 35] |
| $\tilde{\delta}$ | Per cell clearance rate of target cells (and pre-integration) infected cells that facilitate slow integration | 0.01 | [37] |
| $\delta_I$ | Per cell clearance rate of productively infected cells | 0.7 | <sup>b</sup> Inferred from [28, 34-36, 46] |
| $p^j$ | Per cell rate of proliferation of latently infected cells | $p^j - \delta_L^j = 0.0025 \quad j = 0, 1$ | <sup>c</sup> Calculated to ensure that the long-term decay rate of virus in plasma in treated seropositive individuals is given by:<br>$\delta_L^1 + a^1 - p^1 = 5.2 \times 10^{-4}$<br>[9, 10, 30] and assuming<br>$\delta_L^0 = \delta_L^1$ |
| $\delta_L^j$ | Per cell clearance rate of latently infected cells | | |
| $\psi$ | Per cell rate that infected (pre-integration) cells become productively infected | 0.0148 | <sup>d</sup> Calculated from<br>$\psi + \tilde{\delta} = 0.06$ |
| $c$ | Per virion clearance of free virions | 23 days<sup>-1</sup> | [47] |
| $\pi$ | Per cell rate of free virus production | 6000 days<sup>-1</sup> | [28] |
| $z$ | Effectiveness of cART in preventing infection of cells | 0 $\begin{cases} t < 0 \text{ days (post ART)} \\ 1716 \leq t < 1726 \text{ days (post ART)} \end{cases}$<br>0.807 $\begin{cases} 0 \leq t < 1716 \text{ days (post ART)} \\ t \geq 1926 \text{ days (post ART)} \end{cases}$ | Fitted |
| $\phi$ | Effectiveness of antibodies in preventing infection of cells | 0 $t < 1716 \text{ days (post ART)}$<br>0.66 $t \geq 1716 \text{ days (post ART)}$ | Fitted |
| $n$ | Total number of sites | 100 | Chosen to be sufficiently large that saturation of mutations within the genetic region does not measurable impact the accrual of genetic divergence. |
| $\beta$ | Transmission coefficient | $5.253 \times 10^{-10}$ | <sup>d</sup> Fitted to give $R_0=4.6$ [28, 46, 48] |
| $a^j$ | Per cell rate that latently infected cells are activated | $0.003 \text{ days}^{-1}$ $j=0$<br>$7 \times 10^{-6} \text{ days}^{-1}$ $j=1$ | Fitted |
| $\mu$ | Per site probability of substitution upon cell infection | $2 \times 10^{-5}$ | Fitted |
| $k^j$ | Fraction of new infections that result in a latently infected cell | $0.023 \text{ days}^{-1}$ $j=0,1$ | Fitted, assuming $k^0 = k^1$ |

<sup>a</sup> $\delta = 0.15$ is estimated to the mean of two published estimates: $\delta = 0.18$ [28] and $\delta = 0.11$ [35]. <sup>b</sup> $\delta_I = 0.7$ is estimated to the mean of five published estimates: $\delta_I = 1$ [28], $\delta_I = 1$ [36], $\delta_I = 0.38$ [35] and $\mu_I = 0.31$ [46] and $\mu_I = 0.6$ [34]. <sup>c</sup> $\psi + \tilde{\delta} = 0.0248$ is estimated from the decay curves of virus in plasma, described in [8]. <sup>d</sup> The basic reproductive number $R_0 = 4.6$ was estimated by calculating the mean of three published estimates, $R_0 = 1.9$ [28] $R_0 = 3.9$ [46] and $R_0 = 8$ [48]. $R_0$ , is given approximately as: $R_0 = (B\beta\pi)/(\delta\delta_I c)$ and thus $\beta$ was calculated as follows: $\beta = (R_0\delta\delta_I c)/(B\pi)$ .

### Model Equations

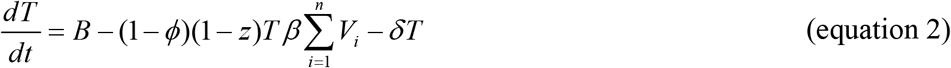

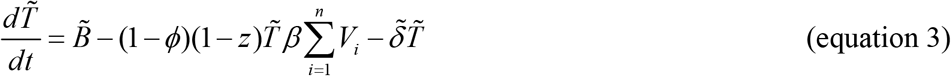

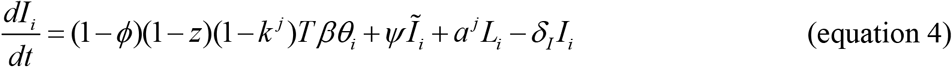

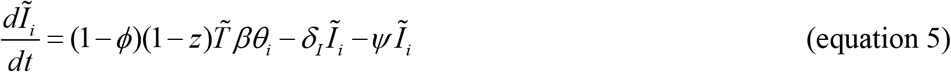

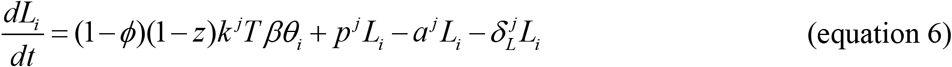

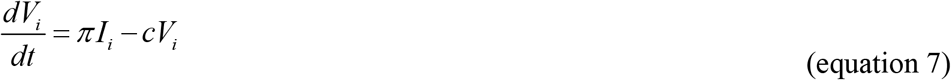

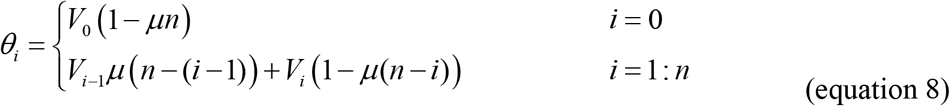

## Acknowledgements

HF, JR and KL were funded by Wellcome Trust and Royal Society grant number 107652/Z/15/Z. TG was funded by the Li Ka Shing foundation.

## Supplementary Information

**Figure S1.**
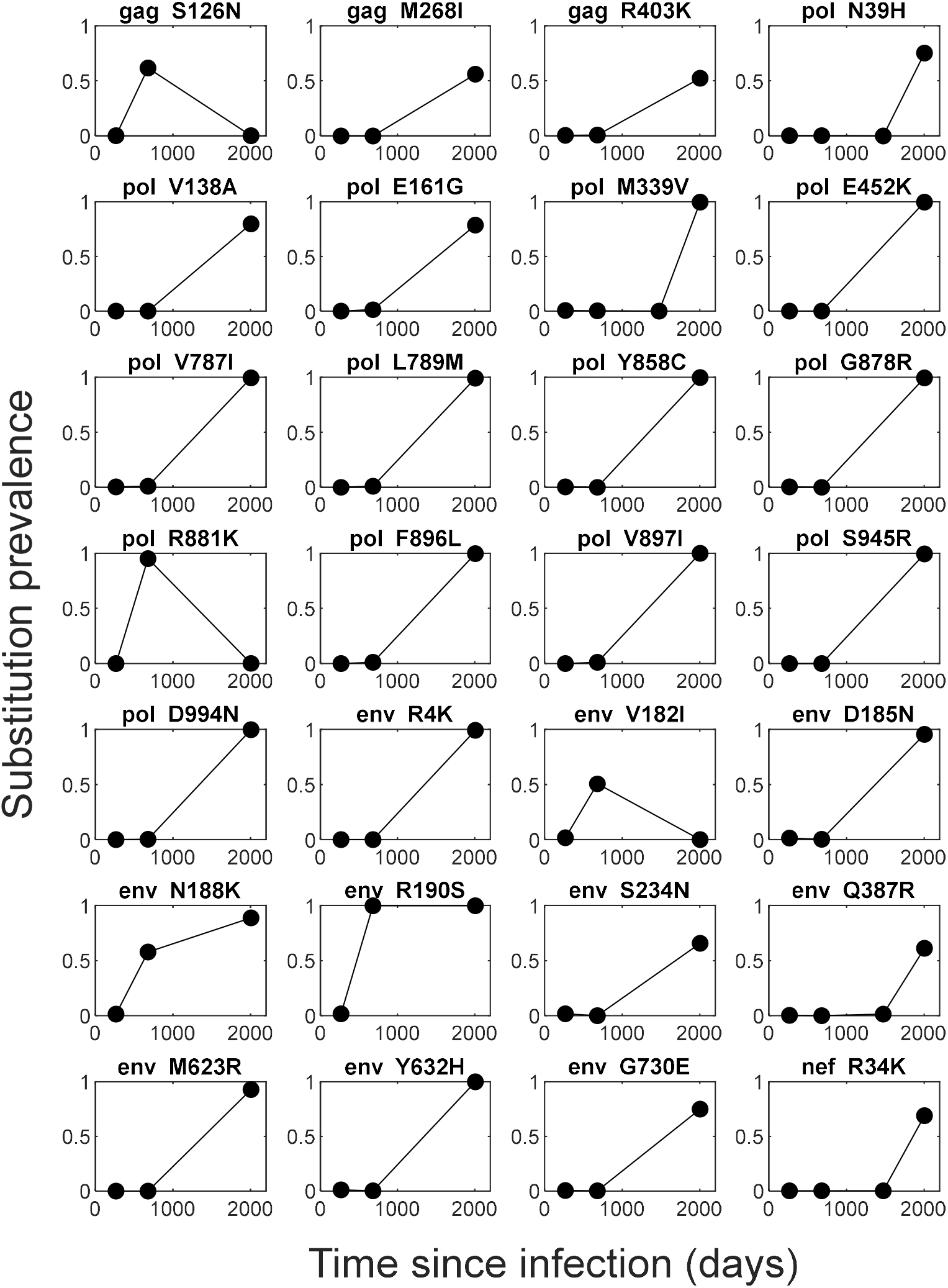
Nonsynonymous sweeps in the seronegative individual.

**Figure S2.**
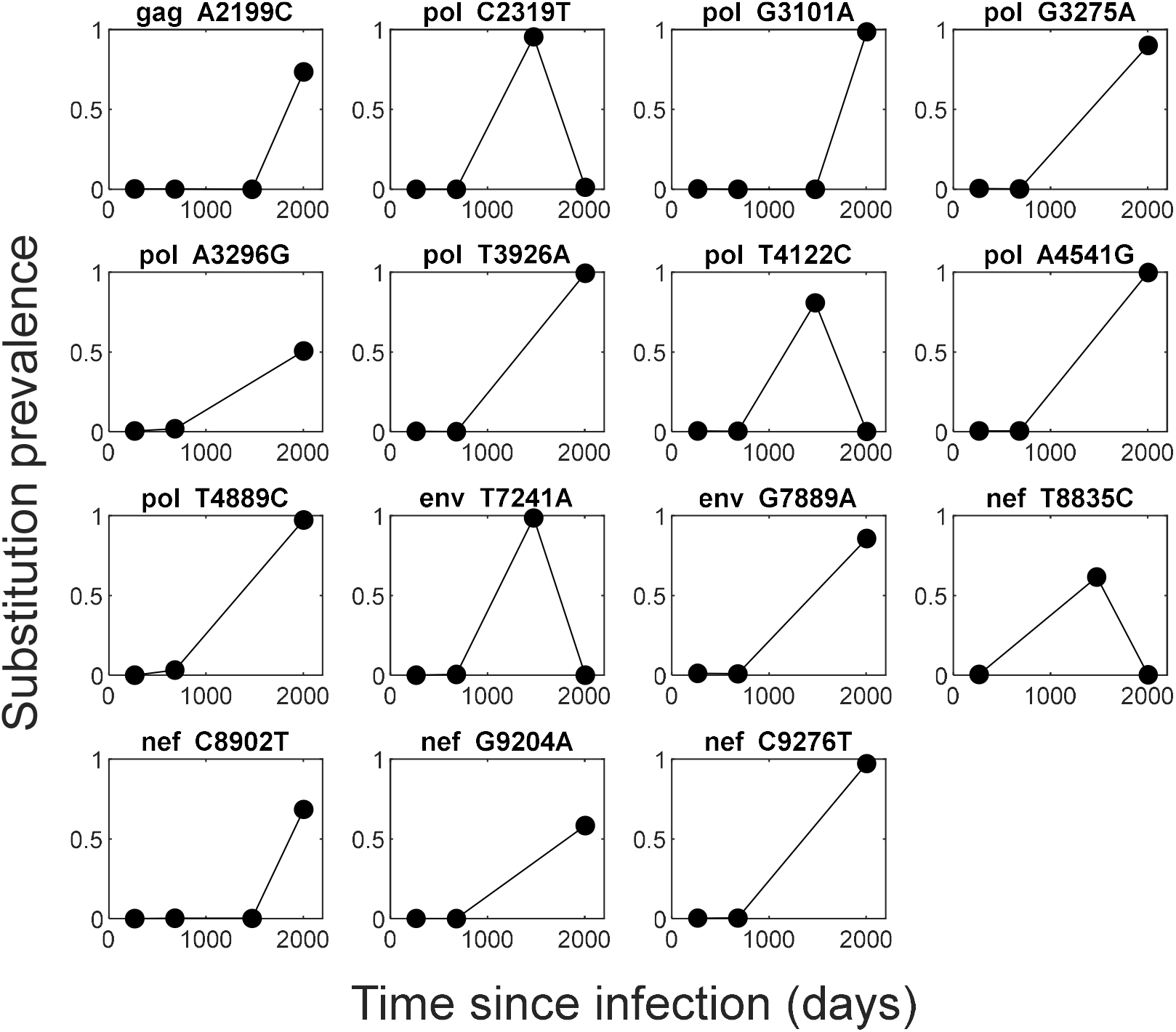
Synonymous sweeps in the seronegative individual.

**Figure S3.**
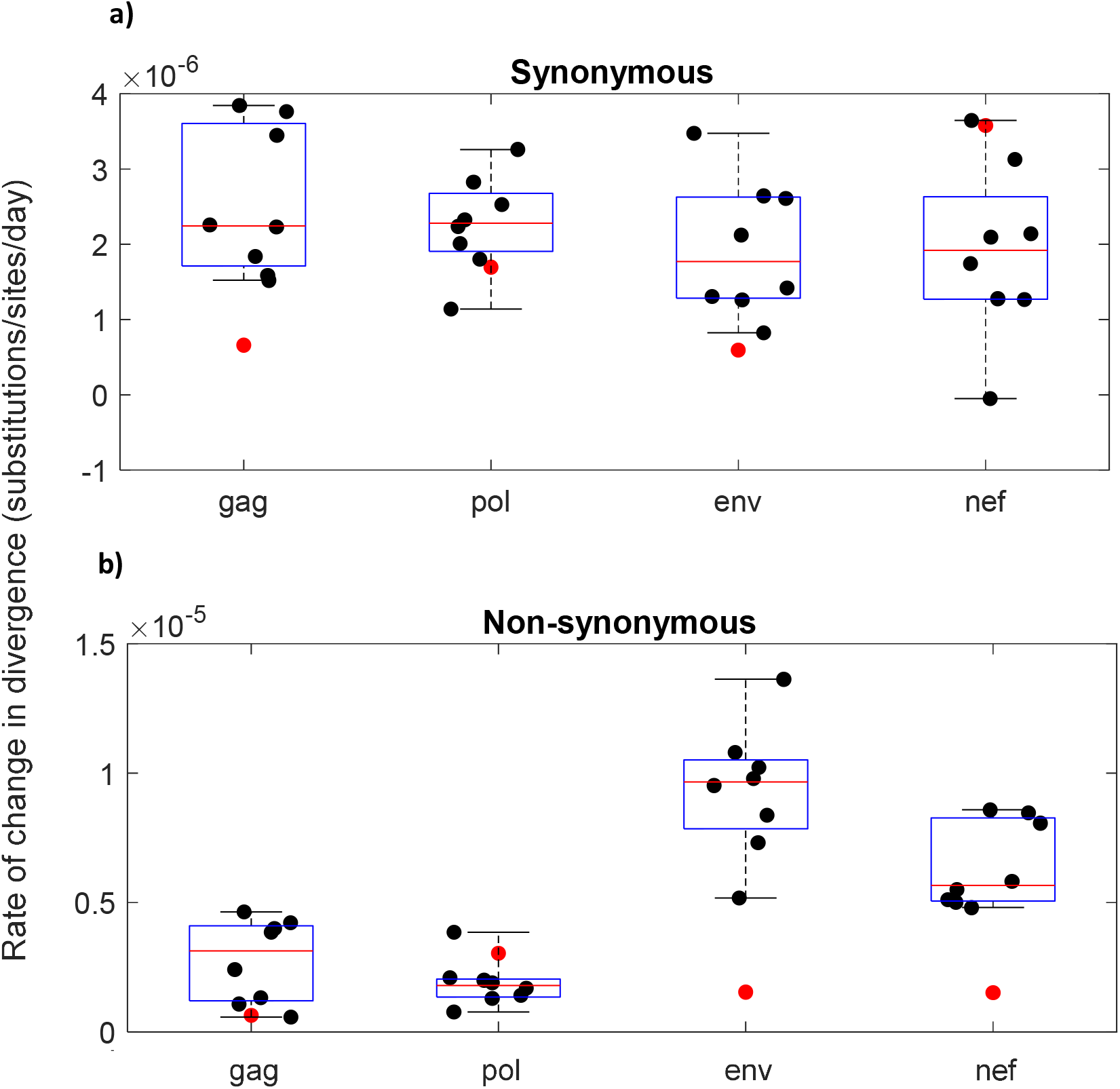
The rate of change in divergence between day 580 and day 2003 post infection in the seronegative patient is compared to the change in divergence over a similar timeframe in the untreated seropositive individuals. The rate of change in synonymous (a) and nonsynonymous divergence (b) between day 580 and day 2003 post infection in the seronegative individual (red markers) is compared to the rate of change in divergence over a similar timeframe in nine untreated seropositive individuals (black markers)[4]. Because the seropositive individuals are not sampled at the same times as the seronegative individual, the change in the divergence in seropositive individuals is estimated using linear regression over the shortest time period spanning the period day 580 to day 2003 post infection. The data are stratified according to gene. The boxplots show the median (red lines), interquartile range (blue box) and full range (black lines) of the rate of change in divergence amongst the seropositive individuals.

**Figure S4.**
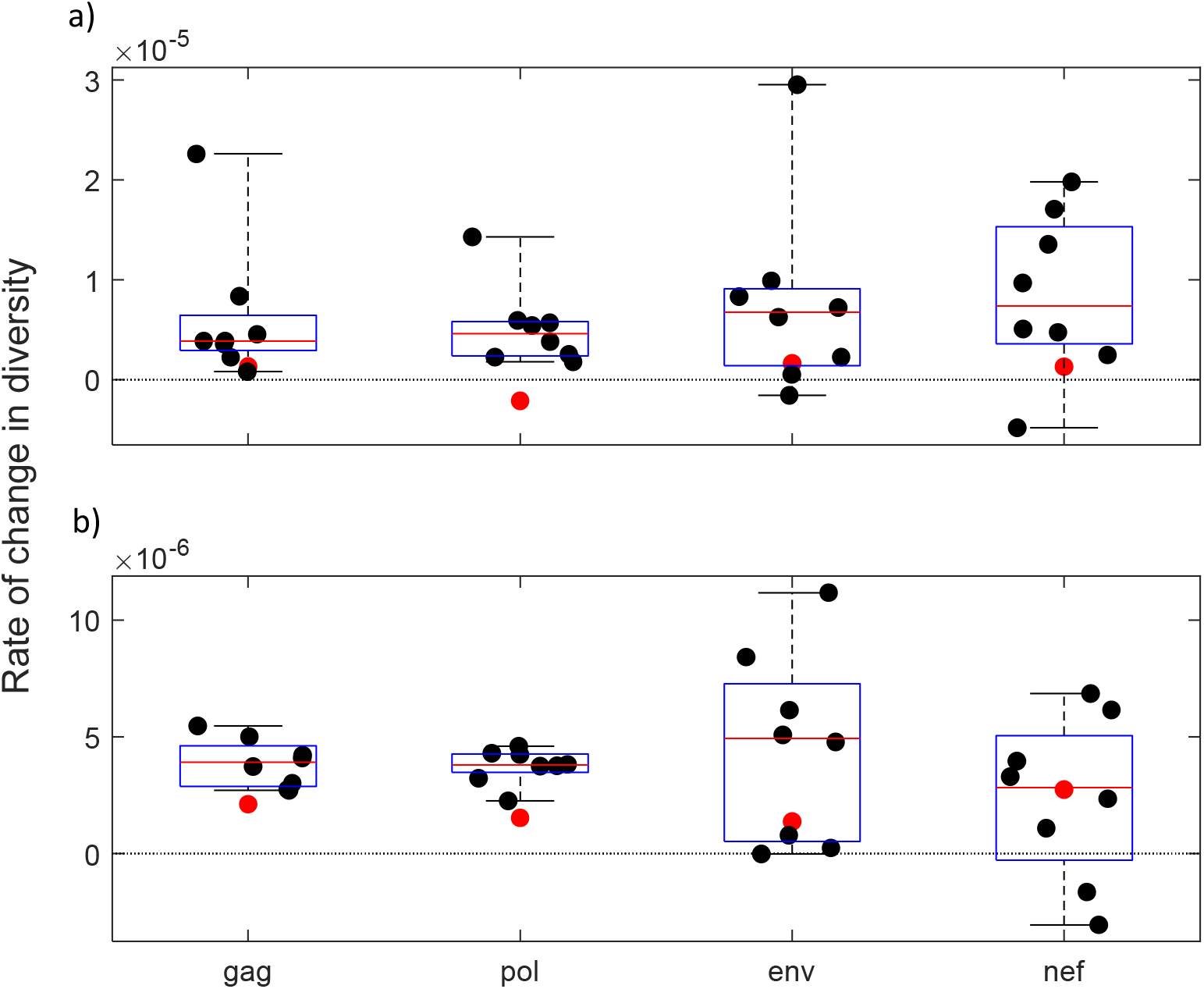
The rate of change in diversity between a) day 266 and day 580 post infection and (b) day 580 and day 2003 post infection is compared to the change in diversity over a similar timeframe in the untreated seropositive individuals. The rate of change in in the seronegative individual (red markers) is compared to the rate of change in diversity over a similar timeframe in nine untreated seropositive individuals (black markers)[4]. Because the seropositive individuals are not sampled at the same times as the seronegative individual, the change in the diversity in seropositive individuals is estimated using linear regression over the shortest time period spanning the corresponding period in the seronegative patient. The data are stratified according to gene. The boxplots show the median (red lines), interquartile range (blue box) and full range (black lines) of the rate of change in divergence amongst the seropositive individuals.

**Table S1.** A summary of read statistics

| Gene | Region length | Days post diagnosis | Number of reads | Mean read length | Mean coverage |
| --- | --- | --- | --- | --- | --- |
| gag | 1512 | 146 | 5099 | 130 | 440 |
|  |  | 560 | 8348 | 137 | 756 |
|  |  | 1355 | 377 | 126 | 31 |
|  |  | 1883 | 9991 | 142 | 935 |
| pol | 3018 | 146 | 11736 | 137 | 536 |
|  |  | 560 | 11762 | 134 | 522 |
|  |  | 1355 | 610 | 131 | 26 |
|  |  | 1883 | 23060 | 151 | 1153 |
| env | 2610 | 146 | 13907 | 142 | 756 |
|  |  | 560 | 21759 | 142 | 1180 |
|  |  | 1355 | 1325 | 129 | 65 |
|  |  | 1883 | 36472 | 146 | 2039 |
| nef | 633 | 146 | 4420 | 122 | 856 |
|  |  | 560 | 5230 | 128 | 1058 |
|  |  | 1355 | 735 | 120 | 140 |
|  |  | 1883 | 9510 | 142 | 2135 |

**Table S2.**
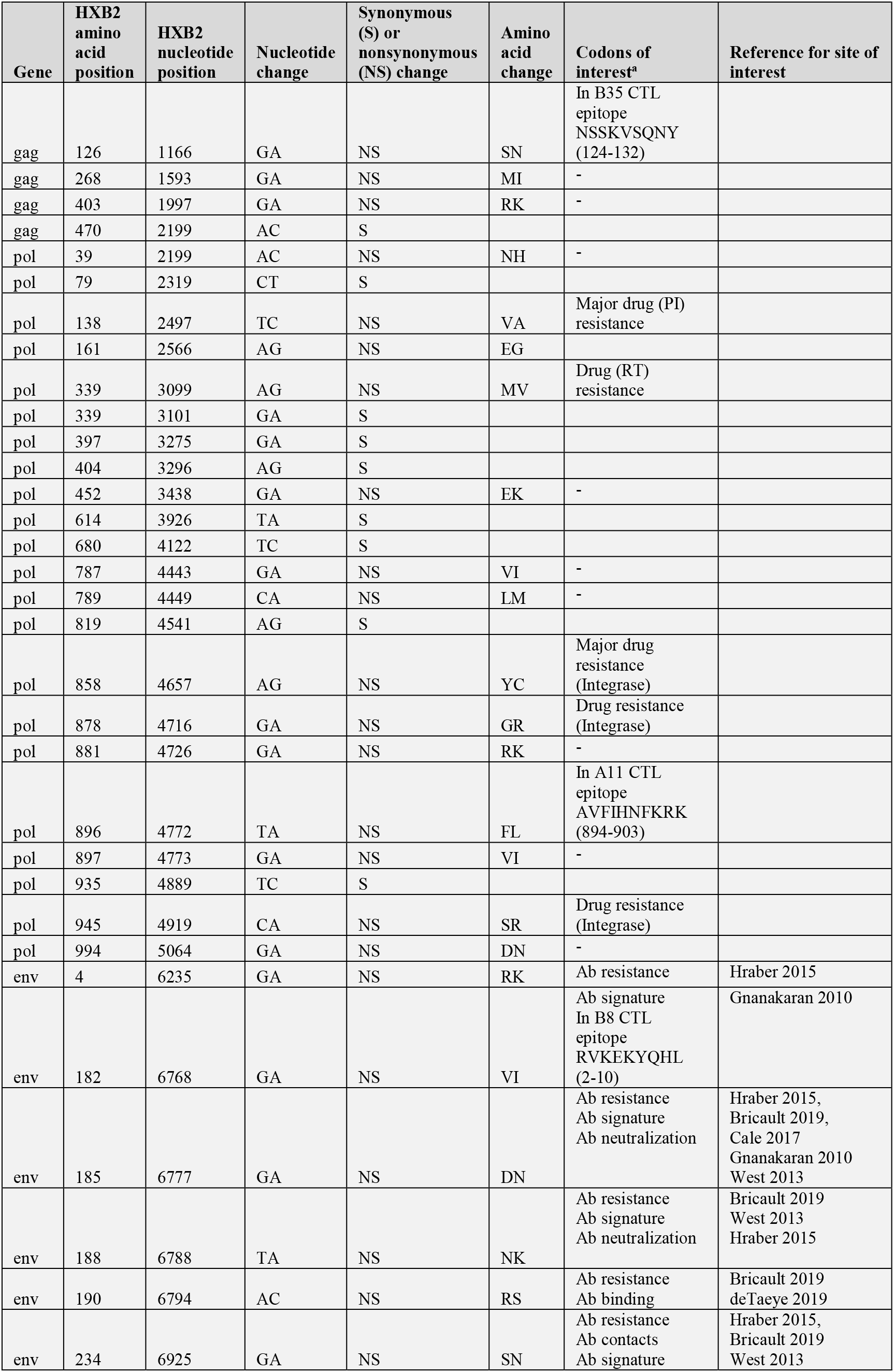
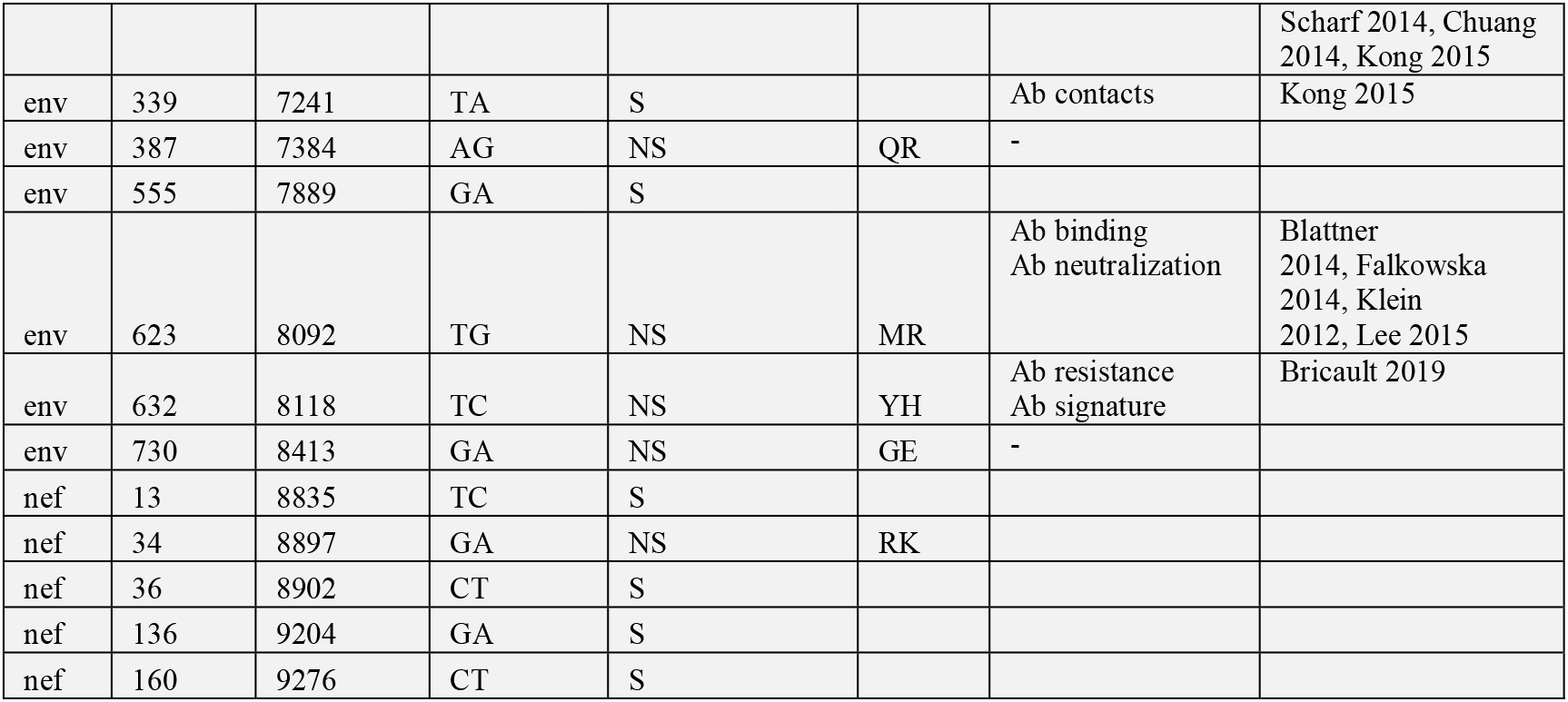
Details of synonymous and nonsynonymous sweeps in the seronegative individual. Sweeps are defined at sites where the prevalence of substitution away from the consensus nucleotide sequence at day 266 post infection (day 146 post cART) is 50% or greater at one or more later sample time. For each of the nonsynonymous sweeps, codons that are associated with described antibody ‘contacts and features’ (https://www.hiv.lanl.gov/components/sequence/HIV/featuredb/search/env_ab_search_pub.comp) or that lie within a ‘best-defined’ CTL epitope (https://www.hiv.lanl.gov/content/immunology/tables/optimal_ctl_summary.html) restricted by the patient’s HLA are listed. Mutational changes were also compared to previously described drug resistance mutations and CTL escape mutations (https://www.hiv.lanl.gov/content/immunology/variants/ctl_variant.html) associated with the patient’s restricting human leukocyte antigen (HLA) type: A*11,*24; B*08, *35; C*07, *04; DRB1*03, *07; DRB3*01; DRB4*Present; and DQB1*02:01, 02. No such CTL escape mutations were identified.

